# Insights into the genetic basis of predator-induced response in *Daphnia galeata*

**DOI:** 10.1101/503904

**Authors:** Verena Tams, Jana Helene Nickel, Anne Ehring, Mathilde Cordellier

## Abstract

Phenotypic plastic responses allow organisms to rapidly adjust when facing environmental challenges - these responses comprise morphological, behavioral but also life-history changes. Alteration of life-history traits when exposed to predation risk have been reported often in the ecological and genomic model organism *Daphnia*. However, the molecular basis of this response is not well understood, especially in the context of fish predation. Here, we characterized the transcriptional profiles of two *Daphnia galeata* clonal lines with opposed life histories when exposed to fish kairomones. First, we conducted a differential gene expression, identifying a total of 125 candidate transcripts involved in the predator-induced response, uncovering substantial intra-specific variation. Second, we applied a gene co-expression network analysis to find clusters of tightly linked transcripts revealing the functional relations of transcripts underlying the predator-induced response. Our results showed that transcripts involved in remodeling of the cuticle, growth and digestion correlated with the response to environmental change in *D. galeata*. Furthermore, we used an orthology-based approach to gain functional information for transcripts lacking gene ontology (GO) information, as well as insights into the evolutionary conservation of transcripts. We could show that our candidate transcripts have orthologs in other *Daphnia* species but almost none in other arthropods. The unique combination of methods allowed us to identify candidate transcripts, their putative functions and evolutionary history associated with predator-induced responses in *Daphnia*. Our study opens up to the question as to whether the same molecular signature is associated fish kairomones-mediated life-history changes in other *Daphnia* species.

## Introduction

Organisms are challenged throughout their lives by environmental changes that have an impact on the health and fitness of each individual. A given phenotype that is advantageous in one environmental setup might become disadvantageous in another. In general, organisms have two possibilities to cope with environmental changes: return to the ecological niche by behavioral (i.e. migration) or physiological changes, or change the boundaries of their ecological niche by genetic adaptation (Van Straalen 2003.). The former is achieved at the phenotypic level and described as a phenotypic plastic response, while the latter is a genetic adaptation process, where genotypes with a higher fitness pass on their alleles to the next generation.

Predation is an important biotic factor structuring whole communities (e.g. Aldana *et al.* 2016; Boaden & Kingsford 2015), maintaining species diversity (e.g. Estes *et al.* 2011; Fine 2015) and driving natural selection in populations (e.g. Kuchta & Svensson 2014; Morgans & Ord 2013). Vertebrate as well as invertebrate aquatic predators release kairomones into the surrounding water (Macháček 1991; Schoeppner & Relyea 2009; Stibor 1992; Stibor & Lüning 1994). In some instances, kairomones can be detected by their prey, inducing highly variable as well as predator-specific responses to reduce their vulnerability. These predator-induced responses are a textbook example of phenotypic plasticity (Tollrian & Harvell 1999) and have been reported in detail for a variety of *Daphnia* species (e.g. Boeing *et al.* 2006; Herzog *et al.* 2016; Weider & Pijanowska 1993; Yin *et al.* 2011).

*Daphnia* are small branchiopod crustaceans and are a model organism widely used in ecology, evolution and ecotoxicology (e.g. Lampert 2011; Miner *et al.* 2012; Picado *et al.* 2007). Members of this genus link trophic levels from primary producers to consumers in freshwater ecosystems and are, therefore, vulnerable to high predation risk (Lampert 2011). Extensive shifts in behavior, morphology and life history traits have been observed in response to predation and predation risk. The responses induced by invertebrate predators include morphological changes such as the formation of helmets in *D. cucullata* (Agrawal *et al.* 1999) and *D. longispina* (Brett 1992) and the formation of neck teeth in *D. pulex* (Tollrian 1995). Vertebrate predators cues have been shown to induce behavioral changes linked to diel vertical migration (Cousyn *et al.* 2001; Effertz & von Elert 2017; Hahn *et al.* 2019) as well as changes in life history traits (Boersma *et al.* 1998; Effertz & von Elert 2017) in *D. magna.* The specificity of such predator-induced responses by vertebrate and invertebrate kairomones has been shown, e.g. for the *D. longispina* species complex from the Swiss lake Greifensee (Wolinska *et al.* 2007). The documented changes in life history traits included a decrease in size at maturity when exposed to fish kairomones and an increase when exposed to kairomones of the phantom midge larvae, a predatory invertebrate of the genus *Chaoborus*.

Although phenotypic plastic responses to predation risk have been extensively studied in the ecological and genomic model organism *Daphnia*, their genetic basis is not well understood (Mitchell *et al.* 2017). Linking predator-induced responses to the underlying genome-wide expression patterns has been attempted from different perspectives (length of exposure time, species and experimental conditions) in *Daphnia*. Orsini *et al.* (2018) investigated the effect of short-term exposure to fish kairomones (several hours) in *D. magna,* revealing no change in gene expression. Yet another study identified over 200 differentially expressed genes in response to invertebrate predation risk in *D. pulex*, of which the most prominent classes of upregulated genes included cuticle genes, zinc-metalloproteinases and vitellogenin genes (Rozenberg *et al.* 2015). Finally, a study on *D. ambigua* under vertebrate predation risk revealed ~50 responsive genes involved in reproduction, digestion and exoskeleton structure (Hales *et al.* 2017).

Our goal is to investigate the genetic basis of life history shifts in response to vertebrate predation risk. *Daphnia galeata* is the ideal candidate species is to address this question, since this species does not show diel vertical migration behavior (Stich & Lampert 1981) or severe morphological changes in the presence of vertebrate predator cues even after long exposure, but diverse shifts in life history traits under vertebrate predation risk (Tams *et al.* 2018). With a combined approach, we aim to understand the complexity of responses to environmental changes such as those induced by predators, which are known to vary across *Daphnia* species. We applied a transcriptomic approach (RNA-sequencing), followed by differential gene expression, gene co-expression and orthology analysis. Gene co-expression network analysis allows to infer gene functions because of the modular structure of co-expressed genes and their functional relations; often co-expressed genes share conserved biological functions (Bergmann *et al.* 2003; Subramanian *et al.* 2005). A further benefit of the co-expression network analysis lies in the opportunity to correlate gene expression and external information (Langfelder & Horvath 2008) thus simplifying the process of candidate genes identification. Additionally, orthology analysis allows revealing functional roles as well as the evolutionary history of transcripts. The degree of conservation of the predator-induced response can be estimated by finding orthologous genes in species having diverged million years ago (Cornetti *et al.* 2019). We hypothesize that a common predator-induced response exists within one *Daphnia* species at the gene expression level and that transcripts involved are evolutionary conserved among *Daphnia* species under vertebrate predation risk.

## Materials and methods

### Experimental organisms

This study was conducted on two *D. galeata* genotypes originally hatched from resting eggs collected from Müggelsee (northeast Germany). A previous study involving 24 genotypes (clonal lines) from four different lakes revealed that the variation for some life history traits increased for genotypes exposed to fish kairomones within the Müggelsee population (Tams *et al.* 2018), meaning a broader range of phenotypes were displayed for that life history trait. We chose the genotypes M6 and M9 which differed in all of their life history traits and were at the contrasting ends of the phenotypic range exhibited by *D. galeata* exposed to fish kairomones. Genotype M6 displayed a phenotype which matured later, produced less offspring and stayed smaller under predation risk. Genotype M9 displayed the opposite phenotype, i.e. matured earlier, produced more offspring and became larger under predation risk (Figure 1).

**Figure 1:**
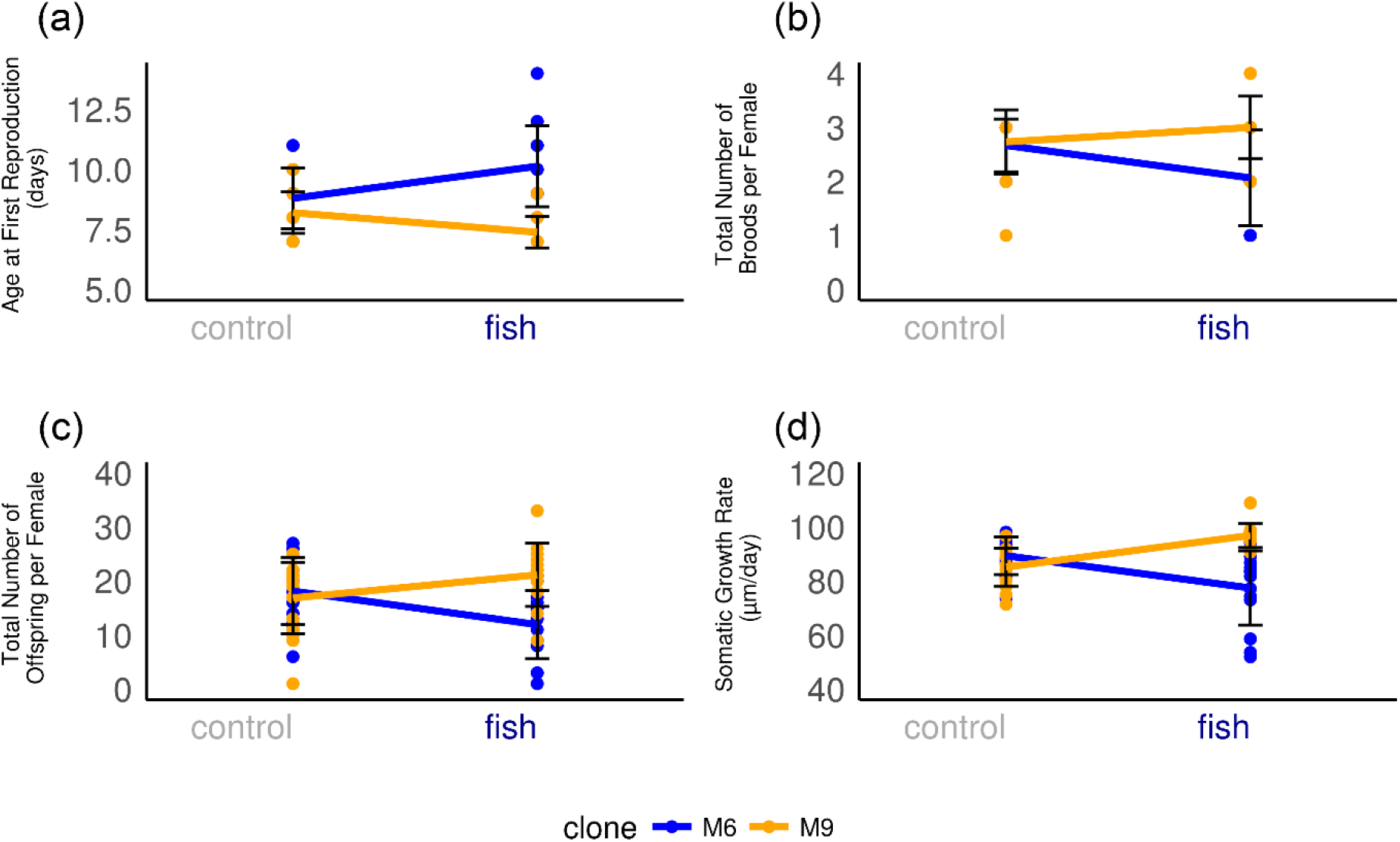
Reaction norms of selected life history traits of experimental genotypes (mean +/− SE) from a previous experiment (Tams *et al* 2018) emphasizing the opposing environmental effect on life history traits for the two genotypes. (a) Age at first reproduction in days. (b) Total number of broods per female (b) total number of offspring per female (d) somatic growth rate in μm per day. ‘Blue’: genotype M6. ‘Orange’: genotype M9. ‘Control’: environment without fish kairomone exposure. ‘Fish’: environment with fish kairomone exposure (predation risk).

### Media preparation

ADaM (Klüttgen *et al.* 1994) was used as the basic medium for fish and *Daphnia* cultures. Two types of media, fish kairomone and control, were prepared and used for breeding and experimental conditions as detailed in Tams *et al.* (2018). Briefly, fish kairomone medium was obtained by maintaining five ide (*Leuciscus idus*) in a 20L tank for 24 hours prior to medium use. All media were filtered (Whatman, membrane filters, ME28, Mixed cellulose-ester, 1.2μm) prior to use and supplemented with 1.0 mg C L-1, P rich *Acutodesmus obliquus*. Media was exchanged daily (1:2) to ensure a nutrient-rich environment and a constant fish kairomone concentration. The algae concentration was calculated from the photometric measurement of the absorbance rate at 800 nm. Cetyl alcohol was used to break the surface tension during breeding and the experiment to reduce juvenile mortality (Desmarais 1997). Breeding and experimental phases were conducted at a temperature of 20°C and a 16h light / 8h dark cycle in a brood chamber with 30% of its maximum light intensity (Rumed, Type 3201D).

### Experimental design and procedures

Each genotype was bred in kairomone-free medium (control) and in fish kairomone medium (predation risk) for two subsequent generations before the start of the experiment to minimize inter-individual variances. To this end, 20 egg-bearing females per genotype were randomly selected from mass cultures. From these females of unknown age, neonates (<24h) were collected and raised under experimental conditions in 750 mL beakers at densities of <40 neonates per beaker. They served as grandmothers (F0) for the experimental animals (F2). Based upon previous work (Tams *et al.* 2018), we started the second (F1) generation after 16-20 days to ensure that offspring from the 3rd to 5th brood were used to start the next generation. The third generation of experimental individuals (F2) was started after 18 days. At the start of the experiment, a pair of neonates was introduced in the experimental vessels (50 mL glass tube) to compensate for juvenile mortality. Before the release of the first brood, on day 6, one of the individuals was randomly discarded whenever necessary so that only one individual remained in each vessel. During the 14 days of the experiment, neonates were removed every 24 hours and the number of broods of each experimental female was documented before media renewal. The adult females (F2) were pooled (n=20) and homogenized in RNAmagic (Bio-Budget technologies, Krefeld, Germany). Five biological replicates were produced per experimental condition (environment) and per genotype, resulting in a total of 400 individuals (two genotypes x two environments x 20 individuals x five biological replicates). Two of these replicates were backup in case of a downstream failure, and three were processed for sequencing (see below). The experiment lasted for 14 days for each experimental individual to assess the long-term effect of fish kairomones on gene expression level in *D. galeata*.

### Data collection and analysis

#### RNA isolation and preparation

Appropriate amounts of RNA were not available from single individuals, and hence we used pools of experimental individuals. Similar pooling approaches have been used in other *Daphnia* differential gene expression studies (Hales *et al.* 2017; Herrmann *et al.* 2018; Huylmans *et al.* 2016; Orsini *et al.* 2018; Ravindran *et al.* 2019; Rozenberg *et al.* 2015). Only experimental females bearing eggs were pooled, resulting in a minor difference in age and experimental time as some experimental females had been pooled a day later. The advantage of sampling females in their inter-molt stage (egg-bearing) is to ensure a stable gene expression (Altshuler *et al.* 2015). Total RNA was extracted from pools of 20 egg-bearing adults after homogenizing in RNAmagic (Bio-Budget technologies, Krefeld, Germany) for 5 min with a disposable pestle and a battery-operated homogenizer. Samples were stored at – 80°C until RNA isolation. Chloroform was added to the homogenate before centrifuging in Phasemaker tubes (Thermo Fisher Scientific, Carlsbad, CA, USA) to separate the upper aqueous and lower phenol phase. The upper aqueous phase was transferred into a clean microcentrifuge tube and the RNA precipitated with absolute ethanol. RNA purification and DNAse treatment were conducted following the Direct-zolTM RNA MiniPrep Kit protocol (Zymo Research, Irvine, CA, USA) with slight modifications. Quality and quantity of purified RNA was checked by spectrophotometry using a NanoDrop 2000 (Thermo Fisher Scientific, Wilmington, DE, USA). The RNA integrity was confirmed with the Agilent TapeStation 4200 (Agilent Technologies, Santa Clara, CA, USA). Only samples showing no degradation and RNA Integrity Numbers (RIN) > 7 were used for subsequent steps. Sequencing was performed for 12 samples (two genotypes × two environments × three biological replicates).

### RNA-seq library construction and sequencing

Library construction and sequencing was identical for all samples and was performed by the company Macrogen (Seoul, South Korea). RNA-seq libraries were constructed using Illumina TruSeq library kits. Illumina HiSeq4000 (San Diego, CA, USA) instrument was used for paired-end sequencing with 101-bp read length resulting in 48-79 million reads per library.

### RNA-seq quality control and mapping

The quality of raw reads was checked using FastQC v.0.11.5 (Andrews 2010). Adapter trimming and quality filtering were performed using Trimmomatic v.0.36 (Bolger *et al.* 2014) with the following parameters: ILLUMINACLIP: TruSeq3-PE.fa:2:30:10 TRAILING: 20 SLIDINGWINDOW: 4:15. After trimming, the read quality was checked again with FastQC to control for the successful removal of adapters. The cleaned reads were mapped to the reference transcriptome of *D. galeata* (Huylmans *et al.* 2016) using NextGenMap v.0.5.4 (Sedlazeck *et al.* 2013) with increased sensitivity (--kmer-skip 0–s 0.0). All reads which had an identity < 0.8 and mapped with a residue number < 25 were reported as unmapped. The option ‘strata’ was used to output only the highest mapping scores for any given read and thus the uniquely mapped reads. The quality of filtering and mapping reads was verified with QualiMap v.2.2.1 (Okonechnikov *et al.* 2016). Subsequently, the htseq-count python script implemented in HTSeq v.0.9.1 was used to quantify the number of reads mapped to each transcript (Anders *et al.* 2015).

### Differential gene expression analysis

Differential gene expression analysis was performed in the R environment v.3.4.2 (R Core Team 2017) with the R package *‘DESeq2’* v.1.18.1 (Love *et al.* 2014) implemented in Bioconductor v.3.6 (Gentleman *et al.* 2004). The calculation was based on normalized read counts per environment (control & fish) using negative binomial generalized linear models. Prior to the analysis, all transcripts with a read count lower than 12 across all libraries were excluded. Results were filtered post-hoc by an adjusted p-value (padj < 0.05) (Benjamini & Hochberg 1995) to reduce the false discovery rate (FDR) and filtered for a log2 fold change ≥ 1. Differentially expressed transcripts (DETs) were binned into four groups: <2-fold, 2- to 4-fold, 4- to 6-fold and >6-fold difference in expression. The three biological replicates were checked for homogeneity by principal component analysis (PCA). A differential expression analysis of genes between environments, between genotypes and between environments within each genotype was done. In addition, a two-factor analysis was applied to investigate genotype-environment interactions (GxE). PCA plots were created in R with *‘ggplot2’* v.2.2.1 (Wickham 2009). The web tool jvenn (Bardou *et al.* 2014) was used to visualize the number of shared transcripts between groups.

### Gene co-expression network analysis

Variance-stabilized read counts obtained from the previous ‘*DESeq2’*-analysis were used in the co-expression analysis. First, an automatic, signed weighted, single gene co-expression network was constructed via an adjacency matrix in the R environment v.3.2.3 with the R package *‘WGCNA’* v.1.61 (Langfelder & Horvath 2008). Second, gene co-expression modules – cluster of highly interconnected genes – were identified based on the topological overlap matrices (TOM) with a soft cut-off threshold of 14 in ‘*WGCNA’*. Module eigengenes (ME) – representing the average gene expression of their module – were calculated and used to investigate their relationship with other modules as well as external information (predation risk and genotype). ME-trait correlations were calculated to identify transcripts of interest with correlation values of > 0.5 or < −0.5. Finally, hubgenes – defined as the most interconnected genes per module – were identified to gain insight into the biological role of a gene co-expression module.

### Gene set enrichment analysis (GSEA)

To identify potential function of differentially expressed and co-expressed transcripts, we assigned Gene Ontology (GO) annotations using the reference transcriptome of *D. galeata* (Huylmans *et al.* 2016). To shed light on the biological importance of transcripts of interest, we performed a gene set enrichment analysis in R with the package *’topGO’* v.2.30.0 (Alexa & Rahnenfuhrer 2016). The default algorithm ‘weight01’ was used taking the hierarchy of GO terms into account, which results in fewer false positive results (Alexa & Rahnenfuhrer 2016). Given that, a multiple testing correction after the Fisher’s exact test was not applied (Timmermans *et al.*). GO terms of the three GO categories ‘Molecular Function’ (MF), ‘Biological Process’ (BP) and ‘Cellular Compounds’ (CC) with a p-value < 0.05 were considered significant.

### Orthology analysis

OrthoMCL cluster information from the reference transcriptome of *D. galeata* (Huylmans *et al.* 2016) was used to enhance our understanding of the functional roles of the transcripts of interest and their evolutionary history. ‘OrthoMCL’ is a tool to identify clusters of homologous sequences in multiple species i.e. orthologs. When assuming that orthologs are functionally conserved, known functions of orthologs in one species can be used to assign putative functions to sequences from other species in the same orthologous group (Li *et al.* 2003). These OrthoMCL clusters were originally build based on data for three *Daphnia* species (*D. galeata*, *D. pulex* and *D. magna*) and two insect species (*Drosophila melanogaster* and *Nasonia vitripennis*). Further details about the genome versions and annotations are available in the original publication by (Huylmans *et al.* 2016).

We analyzed the OrthoMCL clusters containing transcripts of interests by counting how many orthologs from other species where comprised in these clusters. Clusters were grouped into different categories: *Daphnia galeata* specific, *Daphnia galeata* plus one of the other *Daphnia* species (*D magna* or *D. pulex*), *Daphnia* specific, and “all-species” for those containing at least one transcript for each of the reference species (three *Daphnia* and two insect species). This allowed to measure how conserved our signal was and whether the response to predator-risk was affecting transcripts specific to this particular species.

## Results

### RNA-seq data quality

RNA samples passed all quality steps before RNA sequencing. All 12 samples were successfully sequenced, resulting in 48.2 to 79.2 million reads of 101 bp length. After trimming and quality control ~90% of trimmed reads were kept for further analysis. An average of 88% of these trimmed reads were uniquely mapped to the *D. galeata* reference transcriptome. After the filtering process, the full dataset used for further analysis comprised a total of 32,903 transcripts.

### Differential gene expression analysis

Before subsequent analysis, all transcripts with a read count lower than 12 across all libraries were excluded. 23,982 transcripts remained for both genotypes: 21,740 transcripts for genotype M6 and 21,813 for genotype M9.

A principal component analysis (PCA) was performed to visualize the grouping of read counts and to help identify batch effects. The first principal component (PC 1) explained 83% of the variance between genotypes, revealing no clear clustering of read counts per environment (Figure 2A). PC 2 explained just 10% of the variance, which seems to be related to the variance between replicates. Separate plots per genotype improved the visualization of replicate and environmental differences (Appendix 1) but did not indicate an evident clustering by environment either.

**Figure 2:**
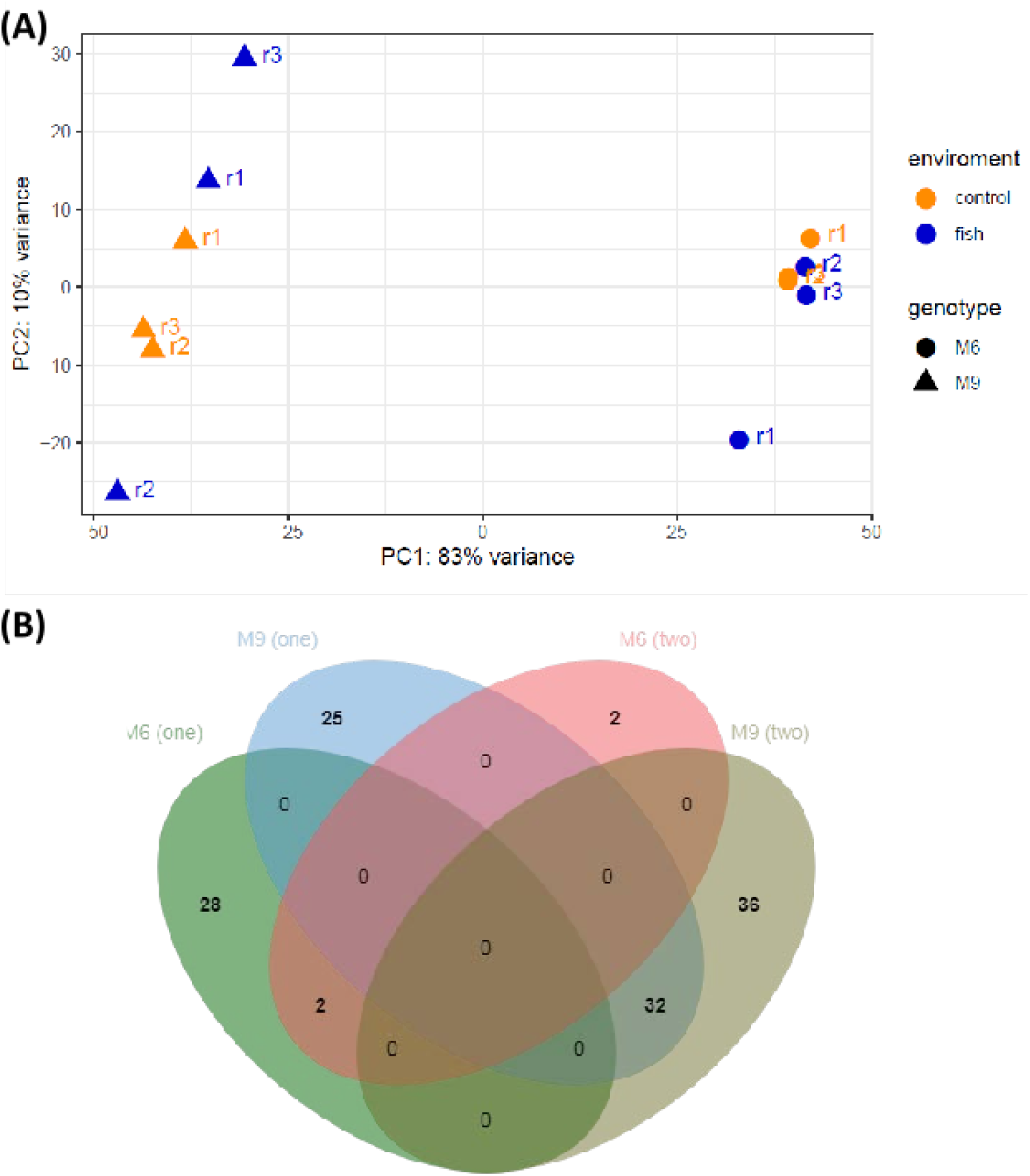
(A) Principal component (PC) plot of the biological RNA-seq samples in *D. galeata*. Yellow: control environment. Blue: fish environment (predation risk). Triangles: genotype M9. Circles: genotype M6. (B) Venn diagram of the 125 differentially expressed transcripts (DETs) related to predation risk in *D. galeata*. The set of DETs originates from the one and two factor analysis. ‘M6 (one)’: DETs from the one-factor analysis for the genotype M6. ‘M9 (one)’: DETs from the one-factor analysis for the genotype M9. ‘M6 (two)’: DETs from the two-factor analysis for the genotype M6. ‘M9 (two)’: DETs from the two-factor analysis for the genotype M9.

The differential expression analysis revealed no differentially expressed transcripts (DETs) between environmental groups, but a total of 5,283 DETs between genotypes (up: 2,228 (42%), down: 3,055 (58%); Figure 3). Because of the strong genotype effect, the genotypes were analyzed separately in a one-factor analysis (Table 1A). For genotype M6, there were 30 DETs between environments (up: 3 (10%), down: 27 (90%)). For genotype M9, there were 57 DETs between environments (up: 21 (37%), down: 36 (63%)). A two-factor analysis accounted for the genotype-environment interaction (GxE) (Table 1B). Between environments genotype M6 had four DETs (up: 1 (25%), down: 3 (75%)) and genotype M9 had 68 DETs (up: 29 (43%), down: 39 (57%)). The GxE resulted in 22 DETs (up: 7 (32%), down: 15 (68%)).

**Figure 3:**
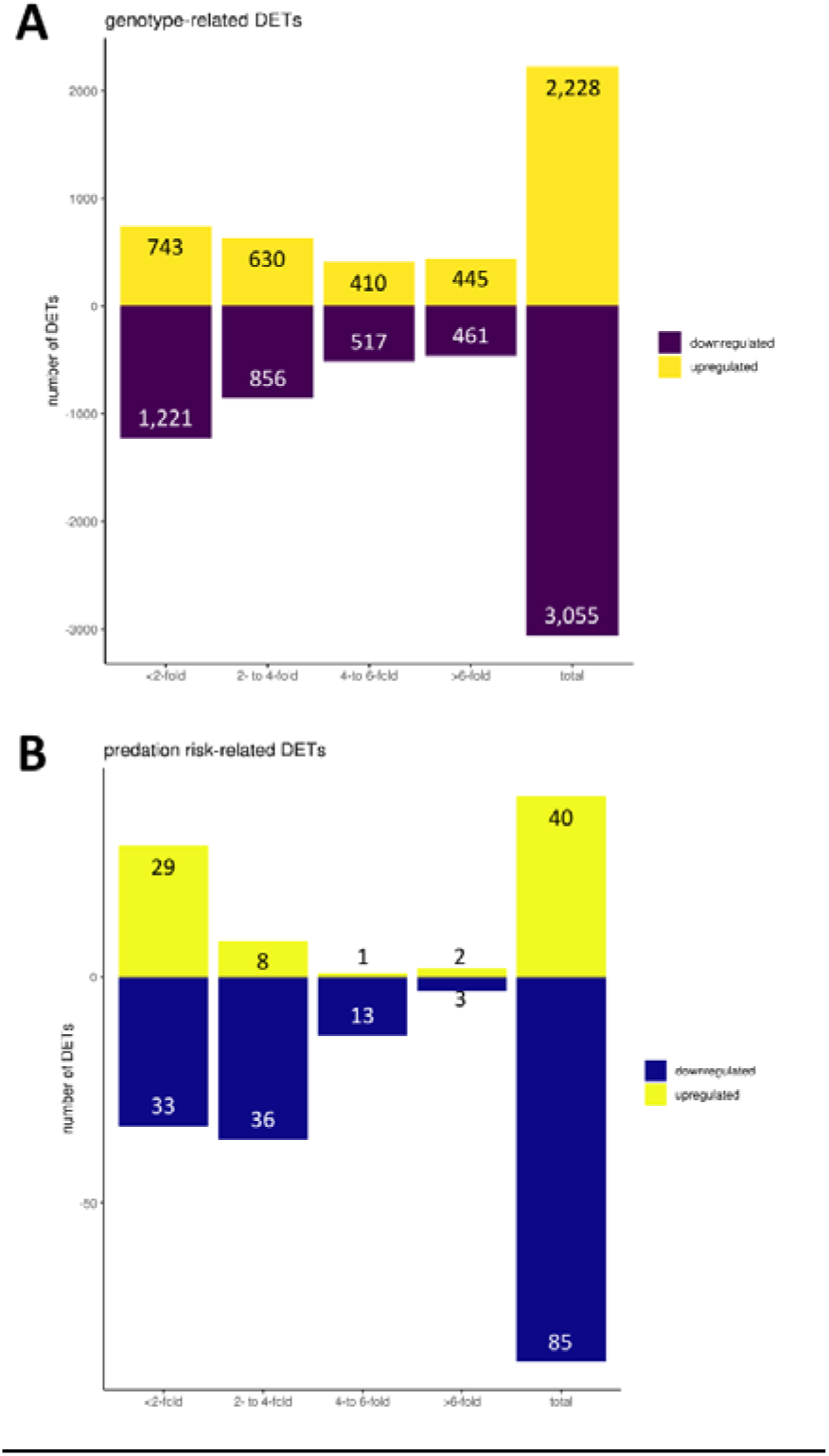
Up- and downregulated differentially expressed transcripts (DETs) grouped by expression foldchange. **(A)** DETs related to genotype. **(B)** DETs related to predation risk.

**Table 1.** Number of differentially expressed transcripts (DETs) in *D. galeata* (p.adj=0.05, foldchange= log2). **(A)** Results of the one-factor analysis. ‘Genotype’: DETs between genotypes (M6 over M9). ‘M6’: DETs within genotype M6 between environments (fish over control). ‘M9’: DETs within genotype M9 between environments (fish over control). **(B)** Results of the two-factor analysis. ‘M6’: environment effect for genotype M6 (fish over control). ‘M9’: environment effect for genotype M9 (fish over control). ‘M6 vs M9’: differences between the two genotypes in control environment (M6 over M9). ‘M6 vs M9 FK’: differences between genotypes in fish environment (M6 over M9). ‘GxE’: genotype-environment interaction (genotype × predation risk).

|  |  |  |  |  |  |  |
| --- | --- | --- | --- | --- | --- | --- |
| **A** |  | All | <2-fold | 2- to 4-fold | 4- to 6-fold | > 6-fold |
|  | Genotype | 5283 | 1964 | 1486 | 927 | 906 |
|  | up | 2228 | 743 | 630 | 410 | 445 |
|  | down | 3055 | 1221 | 856 | 517 | 461 |
|  | M6 | 30 | 11 | 11 | 6 | 2 |
|  | up | 3 | 3 | 0 | 0 | 0 |
|  | down | 27 | 8 | 11 | 6 | 2 |
|  | M9 | 57 | 24 | 27 | 5 | 1 |
|  | up | 21 | 16 | 5 | 0 | 0 |
|  | down | 36 | 8 | 22 | 5 | 1 |
| **B** |  | All | <2-fold | 2- to 4-fold | 4- to 6-fold | > 6-fold |
|  | M6 | 4 | 1 | 2 | 0 | 1 |
|  | up | 1 | 0 | 0 | 0 | 1 |
|  | down | 3 | 1 | 2 | 0 | 0 |
|  | M9 | 68 | 45 | 16 | 6 | 6 |
|  | up | 29 | 22 | 5 | 1 | 1 |
|  | down | 39 | 23 | 11 | 5 | 0 |
|  | M6 vs M9 | 4687 | 1624 | 1204 | 899 | 960 |
|  | up | 1990 | 633 | 494 | 405 | 458 |
|  | down | 2697 | 991 | 710 | 494 | 502 |
|  | M6 vs M9 FK | 3820 | 1114 | 915 | 826 | 965 |
|  | up | 2016 | 611 | 478 | 428 | 499 |
|  | down | 1804 | 503 | 437 | 398 | 466 |
|  | GxE | 22 | 11 | 6 | 4 | 1 |
|  | up | 7 | 3 | 4 | 0 | 0 |
|  | down | 15 | 8 | 2 | 4 | 1 |

No DETs were shared between the two genotypes under predation risk; DETs were shared between the one and two factor analysis only within one genotype (Figure 2B). In total 125 transcripts were differentially expressed between the two environments (hereafter, predation risk-related DETs) (up: 40 (32%), down: 85 (68%); Figure 3, Appendix 2). The differential expression was strong (fold change >2) for downregulated DETs (~60% for predation risk and genotype-related DETs) and for ~67% of upregulated the genotype-related DETs (Table 1, Figure 3). Only ~28% of upregulated, predation risk-related DETs were strongly differentially expressed.

### Gene co-expression network analysis

The single network analysis clustered the expressed transcripts into 16 gene co-expression modules (Figure 4, Table 2). A total of eight modules correlated to predation risk or genotype with a correlation coefficient >0.5 or < −0.5 (Appendix 3). Three small gene co-expression modules associated with predation risk: ‘salmon’ (n= 107), ‘red’ (n= 519) and ‘tan’ (n= 116). The ‘salmon’ module correlated positively with predation risk (p= 0.01); the ‘red’ and the ‘tan’ module correlated negatively with predation risk (p_red_= 0.03, p_tan_= 0.05). Five gene co-expression modules associated to genotype. The two large co-expression modules ‘turquoise’ (n= 5,154) and ‘brown’ (n= 4,760) correlated positively with genotype (p_turquoise_= 0.05, p_brown_> 0.001), while ‘blue’ (n= 4,868), ‘yellow’ (n= 4,612) and ‘green’ (n= 950) correlated negatively with genotype (p_blue_< 0.001, p_yellow_= 0.04, p_green_= 0.04). A dendrogram of the relationship of all co-expression modules showed that the co-expression modules ‘blue’, ‘green’ and ‘yellow’ related closely to genotype and ‘salmon’ to predation risk (Figure 5).

**Figure 4:**
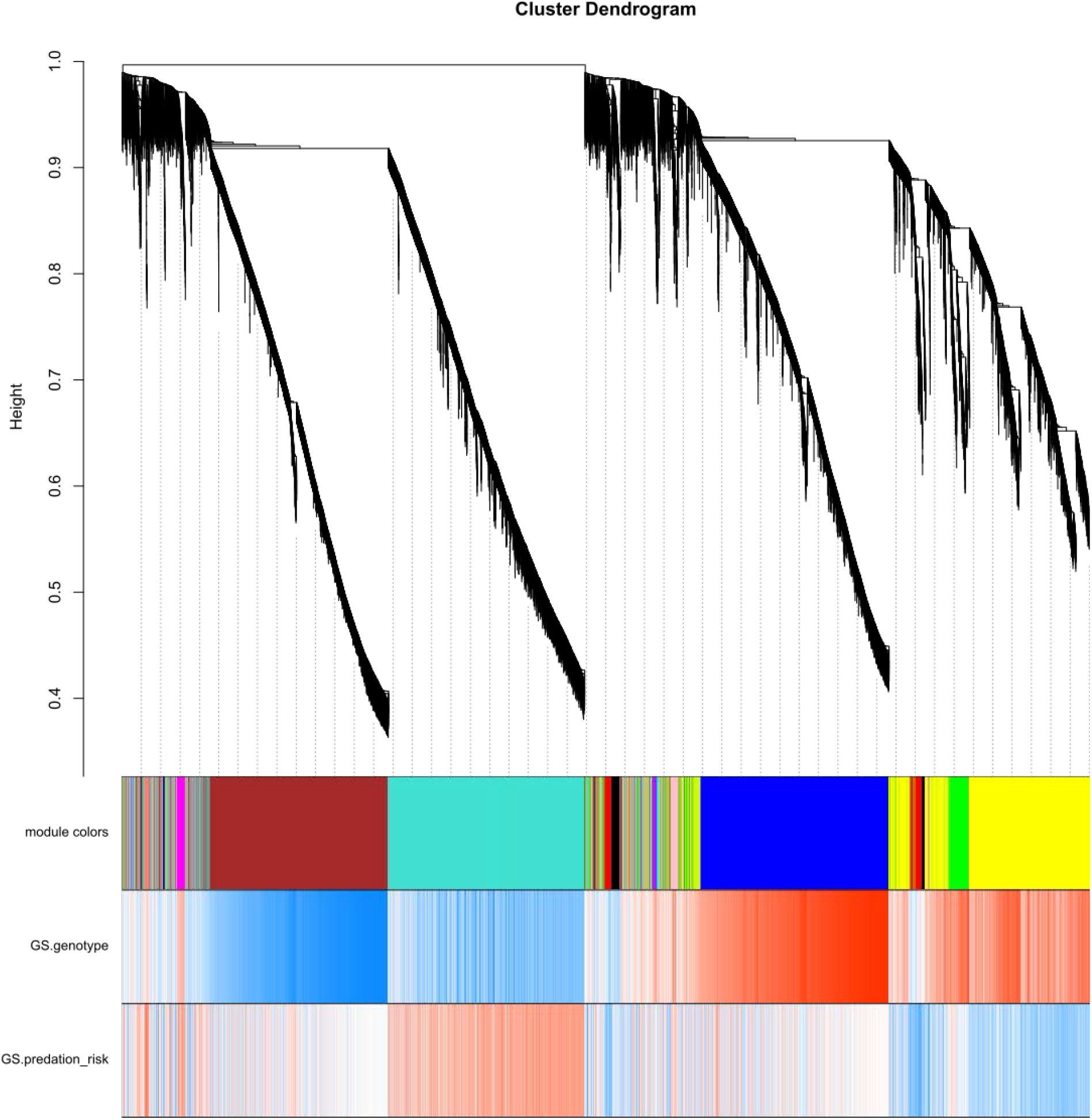
Cluster dendrogram of transcripts in *Daphnia galeata*, with dissimilarity based on the topological overlap matrices (TOM). Additional assignments are module colours, the gene significances (GS) for the trait genotype and predation risk (fish kairomone exposure). ‘Red’: positive correlation of the module with the respective trait. ‘Blue’: negative correlation of the module with the respective trait. Darker hues indicate higher correlation values (darkest hue = 1; lowest hue = 0).

**Table 2.** Overview of gene co-expression modules in *D. galeata*. The table summarizes module color, total number of transcripts per module, the name of the most inter-connected transcript (hubgene), gene significances (GS) and its p-value for predation risk (fish kairomone exposure) and genotype as well as differentially expressed transcripts (DETs) and gene ontology (GO) IDs and classes. The module ‘grey’ contains all co-expressed genes which were not assigned to a co-expression module (n=1,525 (6%)). Significant p-values (p<0.05) are highlighted in bold.

| module | total number of transcripts | hub-gene of co-expression module | GS. predation risk | p.GS predation risk | GS. genotype | p.GS genotype | DETs | GO.ID | GO.class |
| --- | --- | --- | --- | --- | --- | --- | --- | --- | --- |
| turquoise | 5154 | Dgal_a239 | 0.44 | 0.15 | -0.51 | 0.09 | no |  |  |
| blue | 4868 | Dgal_sd37687381411 | -0.02 | 0.95 | 1.0 | **0.00** | no |  |  |
| brown | 4760 | Dgal_s384083 | 0.00 | 0.99 | -1.0 | **0.00** | no |  |  |
| yellow | 4612 | Dgal_o2422d15233t1 | -0.37 | 0.23 | 0.58 | 0.05 | no | GO:0005515 | protein binding |
| green | 950 | Dgal_o7683t4 | 0.06 | 0.86 | 0.57 | 0.05 | no | GO:0042302 | structural constituent of cuticle |
| red | 519 | Dgal_o2656t3 | -0.54 | 0.07 | -0.14 | 0.67 | no | GO:0055114 | oxidation-reduction process |
|  |  |  |  |  |  |  | no | GO:0004497 | monooxygenase activity |
|  |  |  |  |  |  |  | no | GO:0005507 | copper ion binding |
|  |  |  |  |  |  |  | no | GO:0016715 | oxidoreductase activity,..... |
| black | 491 | Dgal_o12661t5 | -0.43 | 0.17 | -0.12 | 0.64 | no | GO:0005515 | protein binding |
| pink | 251 | Dgal_t25528c0t5 | 0.19 | 0.55 | 0.24 | 0.46 | no |  |  |
| magenta | 198 | Dgal_t24643c0t2 | 0.38 | 0.22 | 0.41 | 0.19 | no |  |  |
| purple | 181 | Dgal_o698d42270t2 | 0.02 | 0.95 | 0.50 | 0.10 | no |  |  |
| greenyellow | 127 | Dgal_o21585d23838t2 | 0.04 | 0.89 | 0.61 | **0.04** | no | GO:0005509 | calcium ion binding |
|  |  |  |  |  |  |  | no | GO:0004623 | phospholipase A2 activity |
|  |  |  |  |  |  |  | no | GO:0016042 | lipid catabolic process |
| tan | 116 | Dgal_t6156c0t1 | -0.55 | 0.06 | 0.13 | 0.69 | **yes** |  |  |
| salmon | 107 | Dgal_a 84_f_262622 | 0.65 | **0.02** | 0.03 | 0.93 | **yes** |  |  |
| cyan | 67 | Dgal_t32639c0t1 | -0.15 | 0.64 | 0.29 | 0.36 | no |  |  |
| midnightblue | 56 | Dgal_a 80_j_452081 | -0.43 | 0.16 | -0.04 | 0.91 | **yes** |  |  |
| grey | 1525 | Genes not assigned to a module |  |  |  |  | no |  |  |

**Figure 5:**
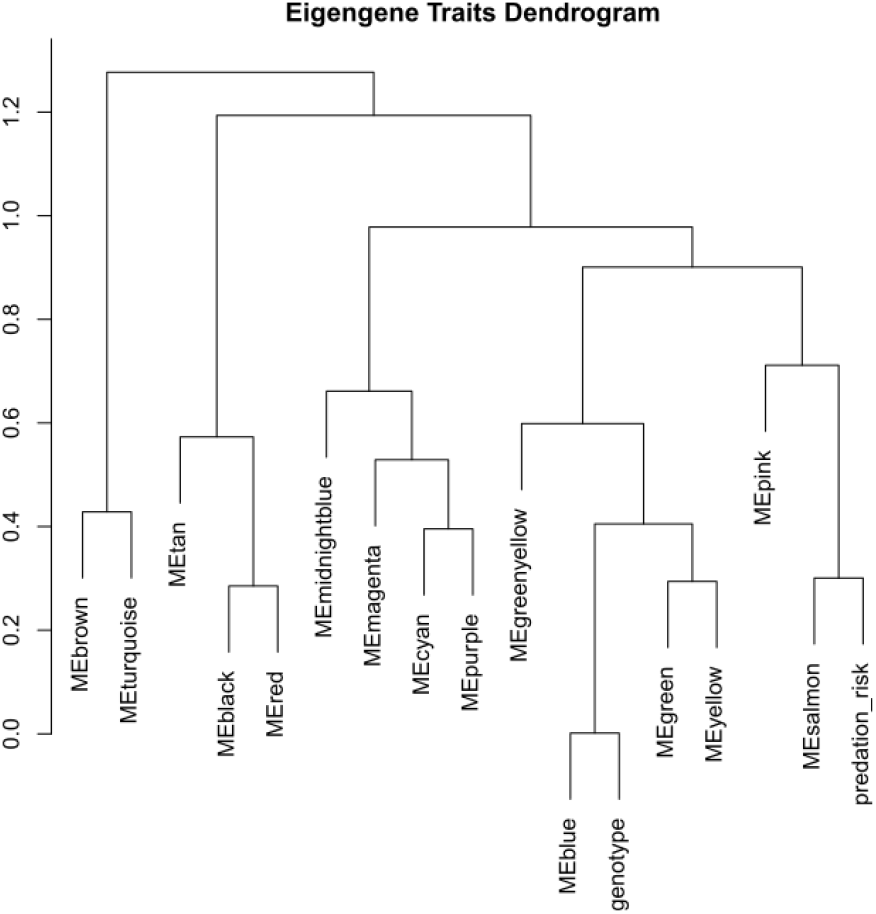
Cluster dendrogram of module eigengenes and external traits genotype and predation risk. This dendrogram shows the relationship between the co-expression modules which are here represented by its module eigengene. In addition, external information genotype and predation risk were included to show their relationship with the gene co-expression modules.

The most highly interconnected gene within a gene co-expression module (hubgene) was identified for each module (Table 2). Three hubgenes of co-expression modules belonged to the previously identified predation risk-related DETs, namely those for the co-expression modules ‘midnightblue’, ‘salmon’ and ‘tan’ (Table 2).

In total 104 of 125 predation risk-related transcripts identified through the differential expression analysis belonged to a co-expression modules of interest (‘salmon’ n=13, ‘tan’ n=9, ‘red’ n=3, ‘turquoise’ n=21, ‘brown’ n=17, ‘blue’ n=7, ‘green’ n=1, ‘yellow’ n=33; Appendix 2).

### Gene ontology (GO) annotation

In total, 10,431 transcripts in the *D. galeata* reference transcriptome had Gene Ontology (GO) annotations (Huylmans *et al.* 2016). The transcript sets of interest were either predation risk- or genotype-related. Predation risk-related transcripts of interest originated from the co-expression modules ‘salmon’, ‘tan’ and ‘red’ (total n= 742), and the differential gene expression analysis (one and two factor analysis; total n= 125). Genotype-related transcripts originated from the co-expression modules ‘turquoise’, ‘blue’, ‘brown’, ‘green’ and ‘yellow’ (total n= 20,344).

36% of transcripts deriving from the co-expression modules of interest were annotated (‘turquoise-blue-brown-green-yellow’ n= 7,117; ‘tan-red-salmon’ n= 207). The lowest rate of annotation (23%) was for genotype-related DETs (n= 1,230 of 5,284) and the highest (33%) for the predation risk-related DETs (n= 41 of 125). Five out of the 15 hubgenes had a GO annotation (Table 2).

### Gene set enrichment analysis (GSEA)

A gene set enrichment analysis was performed on either predation risk-related transcripts of interest (predation risk-related DETs plus transcripts of predation risk-related co-expression modules (‘salmon’, ‘tan’, ‘red’)) or genotype-related (genotype-related DETs plus transcripts of genotype-related co-expression modules (‘turquoise’, ‘blue’, ‘brown’, ‘green’, ‘yellow’)). In total 44 GO terms were significantly enriched in the predation risk-related transcript set; 29 of these GO terms were unique (Appendix 4). In the genotype-related transcript set 209 GO terms were significantly enriched; 168 of them were unique (Appendix 5). There were only eight unique significantly enriched GO terms shared between the predation risk- and genotype-related transcript sets of interest: ‘serine-type endopeptidase activity’ (GO:0004252), ‘extracellular matrix structural constituent’ (GO:0005201), ‘cysteine-type peptidase activity’ (GO:0008234), ‘structural constituent of cuticle’ (GO:0042302), ‘proteolysis’ (GO:0006508), ‘homophilic cell adhesion via plasma membrane adhesion molecules’ (GO:0007156), ‘oxidation-reduction process’ (GO:0055114) and ‘collagen trimer’ (GO:0005581).

### Orthology analysis

GO term and orthoMCL cluster information was available for a total of 9,172 *D. galeata* transcripts. Out of the 867 transcripts in the predation risk-related set 600 were assigned to an orthology cluster. Predation risk-related transcripts of interest were distributed among 563 orthoMCL clusters (2,131 *D. galeata* transcripts; GO annotation n= 224). Most of these orthoMCL clusters comprise orthologs for all three Daphnia species (Figure 6), hinting at a common *Daphnia* response.

**Figure 6:**
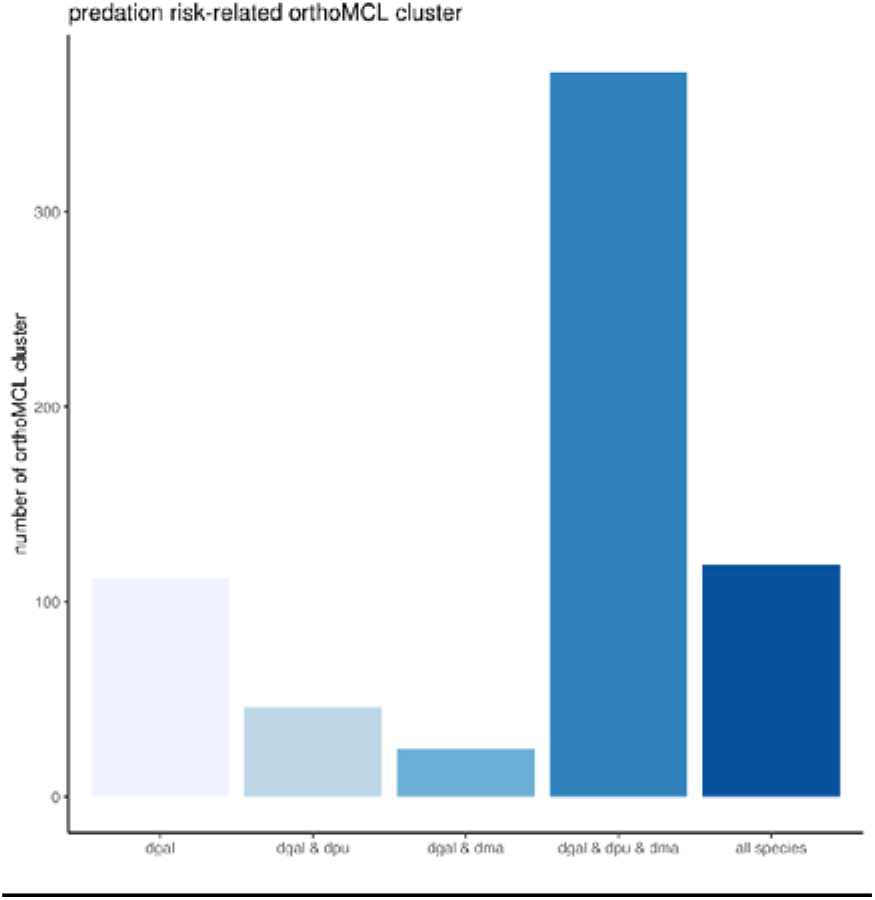
predation risk related orthoMCL clusters grouped according to the species of origin of included transcripts - dgal= *D. galeata*; dma= *D. magna*; dpu = *D. pulex*; all species = all five species included in the analysis.

## Discussion

From an ecological point of view, predator-induced responses in *Daphnia* have been studied extensively. In the past years, few studies addressed the link between such ecological traits and the underlying genetic pathways (Hales *et al.* 2017; Orsini *et al.* 2018; Rozenberg *et al.* 2015). Similar trends in life history shifts after exposure to predator kairomones have been observed across *Daphnia* species showing e.g. the predominant trend of early maturation and a decreased body size under vertebrate predation risk (e.g. Riessen 1999). Thereupon, it seems reasonable to formulate the hypothesis that similar transcripts, differentially expressed, could be involved in the predator-induced response in all *Daphnia* species. To gain insights into the genetic basis of predator-induced responses, we performed gene expression profiling on two *D. galeata* genotypes after long-term exposure to fish kairomones simulating predation risk. We identified a number of transcripts correlating to predation risk and used gene co-expression network analysis, gene ontology annotation and gene set enrichment analysis to describe their putative biological functions. The orthology analysis provided insights into the evolutionary conservation of transcripts, indicating that the majority of transcripts involved in predation risk response were *Daphnia* specific.

### Insights from differential gene expression analysis – similar transcripts, differentially expressed?

In contrast to our expectations, the differential gene expression analysis revealed only a moderate number of differentially expressed transcripts (DETs) between environments within each genotype, but a large divergence between genotypes of the same population in *D. galeata*. Genotype-specific molecular responses to environmental cues were reported for genotypes originating from different populations for *D. pulex* (De Coninck *et al.* 2014) as well as for *D. magna* (Orsini *et al.* 2018). Since we explicitly chose genotypes from one population to minimize the potential genetic variation, population origin can be excluded as an explanation for the observed intraspecific divergence in gene expression profiles in *D. galeata,* concurring with previous studies in our group (Ravindran *et al.* 2019). Instead, the apparent genotype-specific response in our study might be explained by the phenotypic divergence between the two studied clones.

One explanation for the low number of DETs concerning predation risk (environment) compared to genotype-specific differences is that life history changes could only marginally correlate with gene expression. The *D. galeata* genotypes used here only displayed shifts in life history, whereas other *Daphnia* species show additional adaptations of morphology and behavior that could be caused by or correlated to much stronger differential gene expression, e.g. neck-teeth induction that was linked to 230 differentially expressed genes in *D. pulex* (Rozenberg *et al.* 2015).The moderate number of DETs found under predation risk could be explained by other causes than gene expression as well; additional posttranslational processes, such as miRNA-mediated regulation or increased degradation (Schwarzenberger *et al.* 2009) might play a role.

There were three reasons why we expected more pronounced and cumulative changes in differential gene expression in the third experimental generation. First, the chosen *D. galeata* genotypes displayed strong shifts in life history traits after three generations of fish kairomone exposure (Tams *et al.* 2018). Second, the effect of kairomone exposure is expected to be cumulative and to increase over the course of multiple generations; e.g. *D. pulex* displays the largest helmets when exposed to kairomones from *Leptodora kindtii* (an invertebrate predator) for two generations compared to the first generation (Agrawal *et al.* 1999). Third, transgenerational plasticity was described in *D. ambigua* (Hales *et al.* 2017); genes were significantly differentially expressed after one generation of fish kairomone exposure (n=48 DEGs) and without kairomone exposure after the second (n=223 DEGs) and third (n=170 DEGs) generation. To date, it is unknown whether *D. galeata* genotypes display transgenerational plasticity and/or pass on epigenetic modifications after exposure to fish kairomones. Further investigations are therefore required to understand the epigenetic level of inheritance in *Daphnia*.

We expected to find similar transcripts to be involved in the contrasting life history responses of the two genotypes under predation risk. In contrast, a completely different set of transcripts was linked to predation risk within each genotype. The most likely explanation is the high variation in the biological replicates which resulted in no clear distinction between environments. To clarify whether DETs are actually genotype-specific it would be necessary to generate RNA-seq data for more *D. galeata* genotypes from the same and other populations, both with shared and divergent life histories.

### Insights from gene co-expression and gene set enrichment analysis (GSEA) – different transcripts, different functions?

Although different transcripts were identified in the differential expression analysis, the combined approach revealed their similarity of biological functions. In brief, our gene co-expression and gene set enrichment analysis revealed digestion- and growth-related enriched GO terms. These are interesting because predator-induced responses in *Daphnia* include changes in body size (e.g. Tams *et al.* 2018) and morphological modifications (e.g. Laforsch & Tollrian 2004). Such modifications require energy allocated from nutrients; consequently digestive enzymes like peptidases have been shown to be important for juvenile growth rate in *D. magna* (Schwarzenberger *et al.* 2012).

Transcripts of the ‘salmon’ gene co-expression module in *D. galeata* were significantly enriched for ‘serine-type endopeptidase activity’, the most important digestive protease in the gut of *D. magna* (Agrawal *et al.* 2005). The exposure to predator kairomones for one generation in *D. ambigua* – a species from the *D. pulex*-complex more closely related to *D. galeata* than *D. magna* (Adamowicz *et al.* 2009; Cornetti *et al.* 2019) – led to an up-regulation of genes related to digestive functions (Hales *et al.* 2017). Cyanobacterial protease inhibitors cause considerable damage to *Daphnia* populations by inhibiting the gut proteases, impairing their digestion (Schwarzenberger *et al.* 2010). These studies concord with our results suggesting that an increase in ‘serine-type endopeptidase activity’ leads to improved digestion and feeding efficiency that is necessary for the resource allocation that comes with shifts in life history, such as producing a greater number of offspring.

The GO term ‘structural constituent of cuticle’ was significantly enriched in both genotypes suggesting that even though there was no overlap in the affected transcripts, similar functions were affected. The ‘structural constituent of cuticle’ was enriched in *D. pulex* exposed to *Chaoborus* kairomones (An *et al.* 2018; Rozenberg *et al.* 2015) and related to remodeling of the cuticle. Furthermore, it was also enriched in the proteomic response of *D. magna* to *Triops cancriformis* (Otte *et al.* 2015) and is thought to be related to changes in carapace morphology as well as the formation of ultrastructural defenses of the cuticle (Rabus *et al.* 2013). Genes related to body remodeling and activation of cuticle proteins were enriched for *D. magna* exposed to vertebrate and invertebrate predator kairomones (Orsini *et al.* 2018). The investigated *D. galeata* genotypes did not display pronounced morphological defenses, but changes in body size and symmetry especially with regard to head shape (Tams *et al.* 2018). Furthermore, for *D. magna*, *D. pulex* and *D. cucullata*, not only visible morphology changes but also fortification of the carapace in the presence of predator kairomones has been recorded (Laforsch & Tollrian 2004; Rabus *et al.* 2013). Our results indicated that ultrastructural defenses could also be present in *D. galeata*.

Altogether, cuticle-associated proteins seem to play an essential role in the response to vertebrate or invertebrate predators in *Daphnia*. DETs found in genotype M6 showed the possible involvement of ‘metallocarboxypeptidase activity’, known to be involved in the stress response to copper in *D. pulex* (Chain *et al.* 2019).

Interestingly, ‘chitin metabolic process’, ‘proteolysis’, ‘structural constituent of cuticle’, ‘chitin binding’, ‘serine-type endopeptidase’ and ‘metallopeptidase activity’ were all found to be enriched in a gene expression analysis during the molt cycle in the marine copepod *Calanus finmarchicus* (Tarrant *et al.* 2014). Since *Daphnia* need to shed their rigid carapace in order to grow, molting is directly related to changes in body size. Another analysis of *D. magna* exposed to *Triops cancriformis* kairomones revealed the role of proteins related to the cuticle, muscular system, energy metabolism and regulatory proteins that may be involved in morphological carapace defenses and changes in resource allocation (Otte *et al.* 2014). In conclusion, a number of biological functions hypothesized to be involved in kairomone response could be confirmed, e.g., transcripts related to body remodeling and growth.

It is worthwhile to mention that some biologically interesting gene functions were only found with the help of the gene co-expression network analysis and would have been overlooked with only a differential expression analysis. For example, the GO term ‘growth factor activity’ occurred in both ‘red’ and ‘tan’ modules, which correlated negatively with fish kairomone exposure and comprising transcripts not identified as DETs. Nevertheless, they could be extremely important for life history changes and might be directly related to changes in somatic growth rate and body size.

For a more comprehensive understanding of genetic links to phenotypic variation and their biological functions, further annotations and therefore functional tests of candidate transcripts are needed. At present, only one third of transcripts of interest were annotated. When GO annotations progress, a re-analysis might provide new elements for understanding the genetic basis of predator-induced responses in *Daphnia*. Onward, generating gene expression data for all 24 genotypes used in Tams *et al.* (2018) would allow to create models to predict the effect of predation risk for European *D. galeata*. The prediction of reproductive success of *Daphnia* – an ecological keystone species – in response to environmental disturbances or changes is useful to forecast detrimental effects resulting in regime shifts within the ecosystem (Asselman *et al.* 2018).

### Insights from orthology analysis – homologous sequences (common ancestor), evolutionary conserved?

Using orthology – orthologs being homologous sequences that differ because of speciation events – is the most popular strategy to derive functional similarity of sequences (Pearson 2013). Here, we chose this approach to gain insight into the evolutionary conservation of transcripts of interest. The results obtained with the OrthoMCL approach provide support for the hypothesis that phenotypic plastic predator-induced responses might be evolutionary conserved in *Daphnia*.

However, several strategies might have developed over time to cope with or adapt to predation risk. This hypothesis seems likely since *Daphnia* have the ability to rapidly adapt to local predator regimes (Declerck & De Meester 2003) and our study provides elements supporting it. First, the differential gene expression analysis revealed genotype-specific molecular responses to predation risk for genotypes originating from the same population. Second, the involved transcripts have similar functions relating to life history changes induced by predation risk, but different transcripts were involved in the predator-induced response for each genotype. This concurs with the suggestion of niche-specific adaptation in *D. magna* due to the genotype- and condition-specific transcriptional response to environmental changes of biotic and abiotic factors (Orsini *et al.* 2018). Their gene co-expression analysis revealed that genes of interest were crustacean related, meaning that the conservation of genes did not exceed the level of crustaceans (Orsini *et al.* 2018). Further insights into evolutionary conservation of differentially and/or co-expressed transcripts linked to phenotypic traits are available for modern and ancient (resurrected) *D. pulicaria* exposed to different phosphorous regimes (Frisch *et al.* 2020). With a different, yet similar approach this recent study reveals the importance of a holistic approach to tackle the question: What is the molecular basis of phenotypic responses to environmental changes?

Gene expression analyses to uncover the molecular basis of predation-induced phenotypic changes have focused so far on single species, and invertebrate predators. Studies conducted in similar conditions are lacking and prevent us from drawing conclusions about a general *Daphnia* response to fish predation. In the future, simultaneous exposure of several species to kairomones, and the coupling of phenotyping and gene expression would help to address the question of a conserved response.

## Conclusion

In summary, the aim of this study was to characterize the genetic basis for the predator-induced response of the freshwater grazer *D. galeata*. Our hypothesis that clonal lines present a common predator-induced response by regulating the expression of the same transcript set could not be confirmed. However, transcripts with similar biological functions – relating to digestion and growth – were identified for the genotypes with the same population origin under predation risk. The transcriptional profiling revealed differentially expressed transcripts and gene co-expression modules in connection to predator-induced responses in *D. galeata*. The biological functions discovered here represent a valuable starting point for future investigations addressing the functionality of certain transcripts *per se* or in respect to a response to environmental changes. For example, by providing detailed lists of candidate transcripts one can choose specific candidates to test their biological functions in knock-down experiments. Lastly, orthology analysis revealed that predation risk-related transcripts possess orthologs in other *Daphnia* species, suggesting that phenotypic plastic predator-induced responses are evolutionary conserved, and warranting further investigation.

## Data Accessibility Statement

Raw RNA-seq reads for all 12 samples and the experimental set up for the analysis of DETs are available from ArrayExpress (accession E-MTAB-6234). Raw read counts, custom scripts and supplementary tables will be made available on Dryad upon publication.

## Authors’ contributions

The study was designed by VT, JHN and MC; laboratory work was carried out by JHN, AE and VT; gene expression and gene network analysis was done by JHN and VT; VT, JHN and MC wrote the manuscript, all authors gave final approval.

## Acknowledgments

We thank Jonny Schulze for his help during *Daphnia* breeding and the experiment. This work was supported by the Volkswagen Foundation (Grant No. 86030). Animal handling and experiments were in accordance with the ethical standards (approved for the execution of experiments on vertebrates, No. 75/15). We thank Suda Parimala Ravindran for her advice and sharing the annotation information of the *Daphnia galeata* transcriptome. Lastly, we would like to thank Jennifer Lohr for her language check and very useful editing, as well as two anonymous reviewers for comments on an earlier version of this manuscript.

## Appendix

## Appendix 1.

Principal component (PC) plot of the biological RNA-seq samples of *D. galeata.* **(A)** genotype M6 **(B)** genotype M9. ‘Orange’: control environment. ‘Blue’: fish environment (predation risk).

## Appendix 2.

List of all differentially expressed transcripts (DETs) in *D. galeata* in response to fish kairomone (predation risk). The list contains 125 predation risk-related DETs including co-expression module, OrthoMCL cluster information and GO term annotation. Hub-genes are highlighted in bold.

## Appendix 3.

Heatmap of correlation values of module eigengenes and external traits genotype and predation risk. Red and blue indicate a positive and negative correlation of the module with the respective trait. Darker hues indicate higher correlation values.

## Appendix 4.

List of significantly enriched gene ontology (GO) terms for the predation risk-related dataset of *D. galeata*. Only significantly enriched GO terms are shown in ascending order (classicFisher <0.05). GO IDs which occur more than once are highlighted in grey.

## Appendix 5.

List of significantly enriched gene ontology (GO) terms for the genotype-related dataset of *D. galeata*. Only significantly enriched GO terms are shown in ascending order (classicFisher <0.05). GO IDs which occur more than once are highlighted in grey.

