## Supplementary figures and images for "Insights into the genetic basis of predator-induced response in *Daphnia galeata*"

### Appendix 1

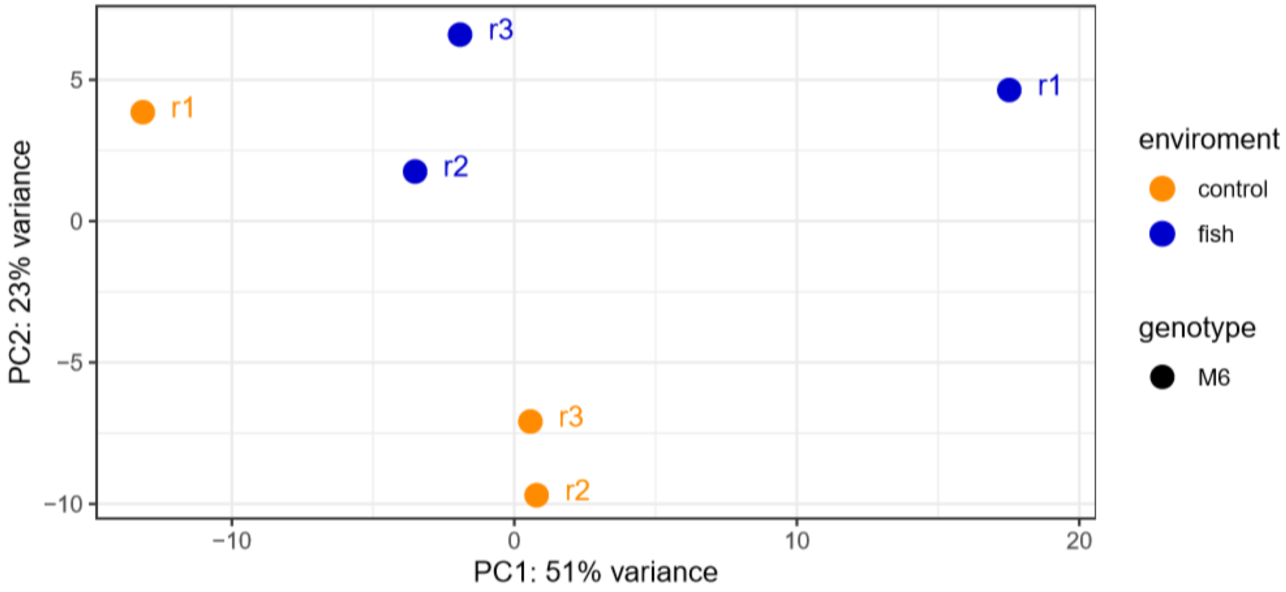

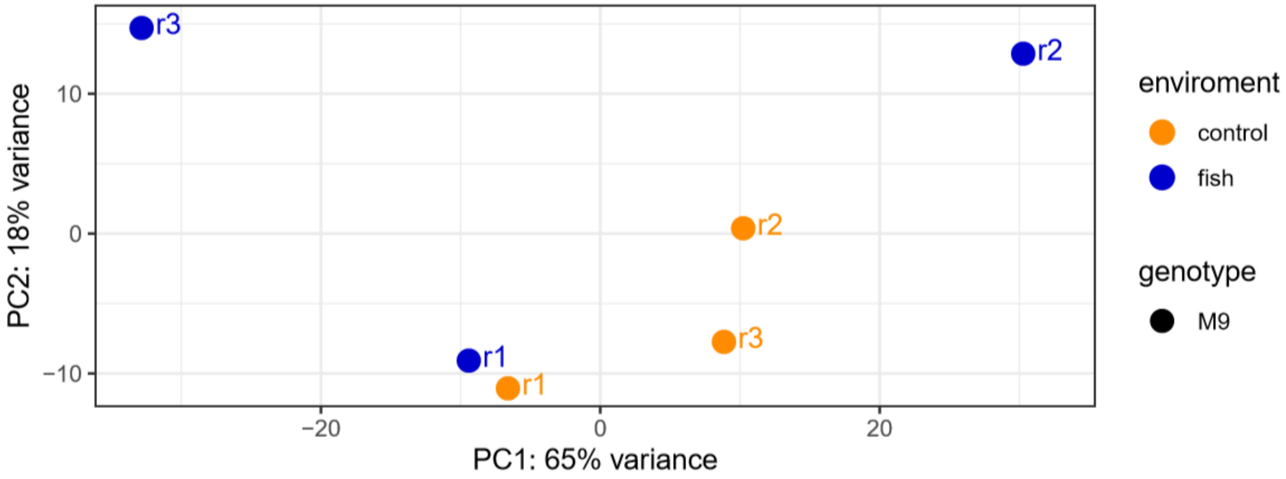

### Appendix 3

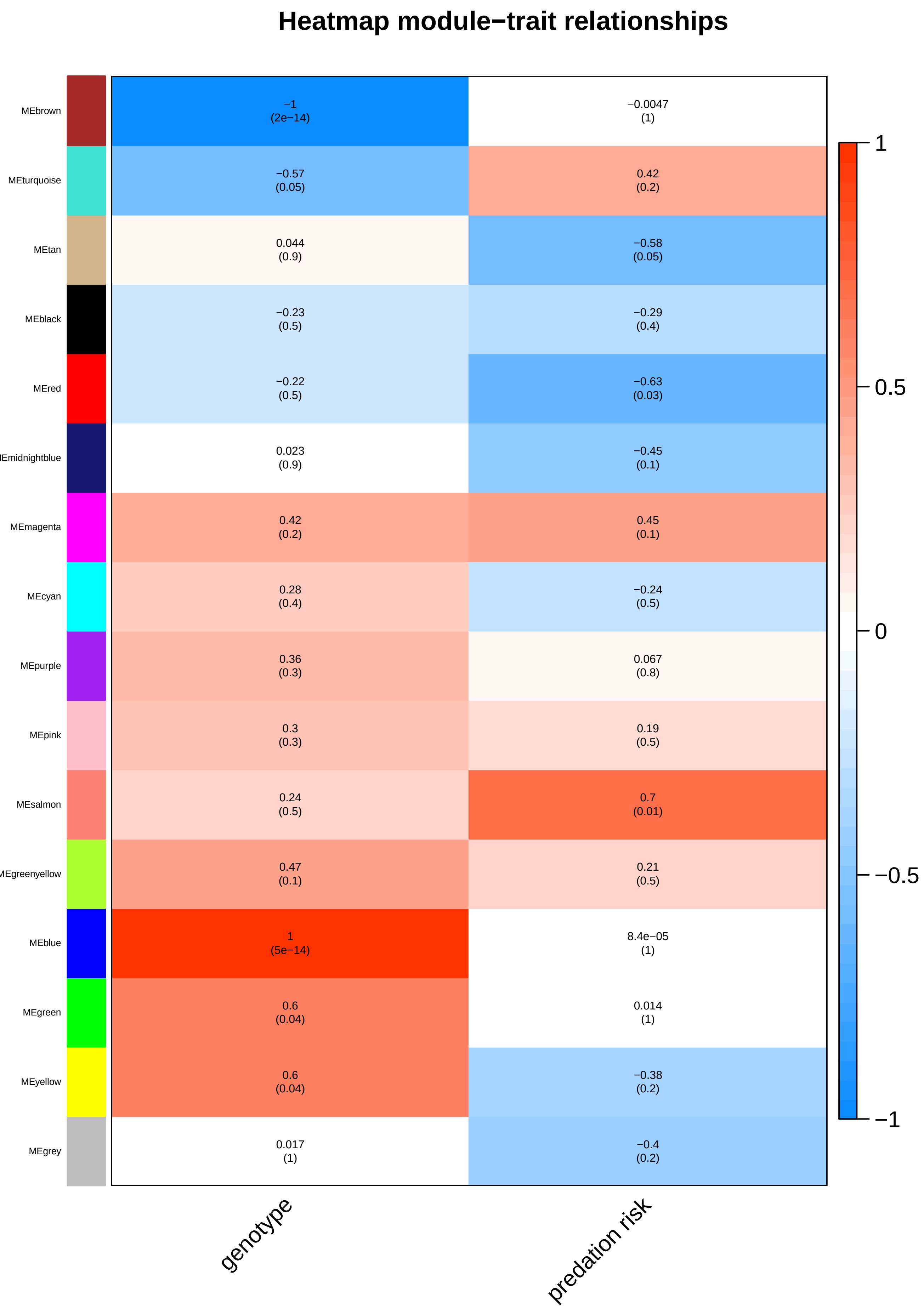
